# Airborne environmental DNA captures terrestrial vertebrate diversity in nature

**DOI:** 10.1101/2022.10.24.512985

**Authors:** Christina Lynggaard, Tobias Guldberg Frøslev, Matthew S. Johnson, Morten Tange Olsen, Kristine Bohmann

## Abstract

The current biodiversity and climate crises highlight the need for efficient tools to monitor terrestrial ecosystems. Here, we provide evidence for the use of airborne eDNA analyses as a novel method to detect terrestrial vertebrate communities in nature. Metabarcoding of 143 airborne eDNA samples collected during three days in Åmosen Nature Park, Denmark yielded 64 bird, mammal, fish and amphibian taxa, representing about a quarter of the around 210 wild terrestrial vertebrates that have been registered in the greater Åmosen area through years of compiling observational data. We provide evidence for the spatial movement and temporal patterns of airborne eDNA and for the influence of weather conditions on vertebrate detections. This study demonstrates airborne eDNA for high-resolution biomonitoring of vertebrates in terrestrial systems and elucidates its potential to guide global nature management and conservation efforts in the ongoing biodiversity crisis.

## Introduction

Vertebrates play important roles in the Earth’s ecosystems, shaping ecosystem structure and functioning through pollination ^1^, seed dispersal ^2^, foraging and predation ^3–5^ and ecosystem engineering ^6^. Further, they can provide ecosystem services ^7–9^, reduce disease transmission ^10^ and serve as indicators of ecosystem health ^11^. However, numerous vertebrates are currently threatened by extinction, population declines and displacement and with that, ecosystem functioning and services are lost and conservation efforts are needed ^12^. To inform and assess nature management and conservation efforts and to provide data for ecosystem and biodiversity studies, effective tools to gather data on species presence in space and time are in great demand ^13^.

In terrestrial ecosystems, the occurrence of vertebrate species in space and time can be mapped using visual observation or capture of individuals, identification of their scats, foot prints and vocalisations ^14,15^, and using camera trapping ^16^. In addition, environmental DNA (eDNA) approaches can be employed to detect terrestrial vertebrates, for example targeting eDNA in freshwater ^17,18^, soil ^19^ or in the guts of parasitic, scavenging or coprophagous invertebrates ^20–23^. However, while effective, the substrates that are currently used to sample the biodiversity of terrestrial vertebrates can be tedious and expensive to collect, and prone to bias towards certain taxonomic groups ^23,24,reviewed in 25^.

Recently, it was demonstrated that terrestrial vertebrates can be detected through sequencing of airborne eDNA ^26,27^. However, while they demonstrated airborne eDNA as an untapped and highly promising source for studying and monitoring terrestrial vertebrate communities, the studies were conducted in urban zoo environments. These pilot studies were highly promising but the efficacy of the airborne eDNA for detection of vertebrates in such environments will most likely differ from natural environments. In zoos, animal species are highly concentrated and confined to the same space over long time spans. Further, there can be transport of bioaerosols and bioaerosol precursors through human activity, e.g. during cleaning of enclosures and stables and when visitors walk in and around the zoo ^26^. In contrast, in natural environments, terrestrial vertebrates will not be as concentrated and will not be confined to the same limited areas. Also, in zoos, buildings can create physical barriers for bioaerosol movement, whereas human movement and disturbance may facilitate dispersal ^26^. In nature, bioaerosol movement might be hindered by dense vegetation and natural physical features, whereas movement of bioaerosols may be facilitated by other species, such as large terrestrial herbivores and less so by humans. Therefore, while the results of the zoo-based studies are promising, it is necessary to determine the applicability of airborne eDNA to obtain terrestrial vertebrate biodiversity data in natural settings to demonstrate the applicability of this technique in nature. Here, we explore the utility of airborne eDNA for monitoring of terrestrial vertebrate communities in a natural environment. By sequencing airborne eDNA collected in a forest in Åmosen Nature Park in western Zealand, Denmark, we aim to i) demonstrate use of airborne eDNA to monitor terrestrial vertebrates in nature, ii) improve our understanding of how to collect airborne eDNA in natural settings, iii) explore the temporal and spatial distributions of airborne eDNA signals.

## Results

We collected air filtering samples in Åmosen Nature Park, Denmark, in the boreal autumn, September-October 2021 (Fig. 1). Samples were collected in two parallel north-south transects, each comprising three sampling sites, and extending from a stream at the forest edge into a mixed deciduous and pine forest. At each sampling site, duplicate sets of low air flow (1.1 m^3^/hr using a 5 V fan) and high air flow (3.5 m^3^/hr using a 12 V fan) samplers were set up, with air filters replaced approximately every 12 hours for three consecutive days and nights, resulting in a total of 144 air filtering samples. To ensure that samples represented airborne eDNA, one filter was discarded from the analyses as it was exposed to rain.

**Figure 1.**
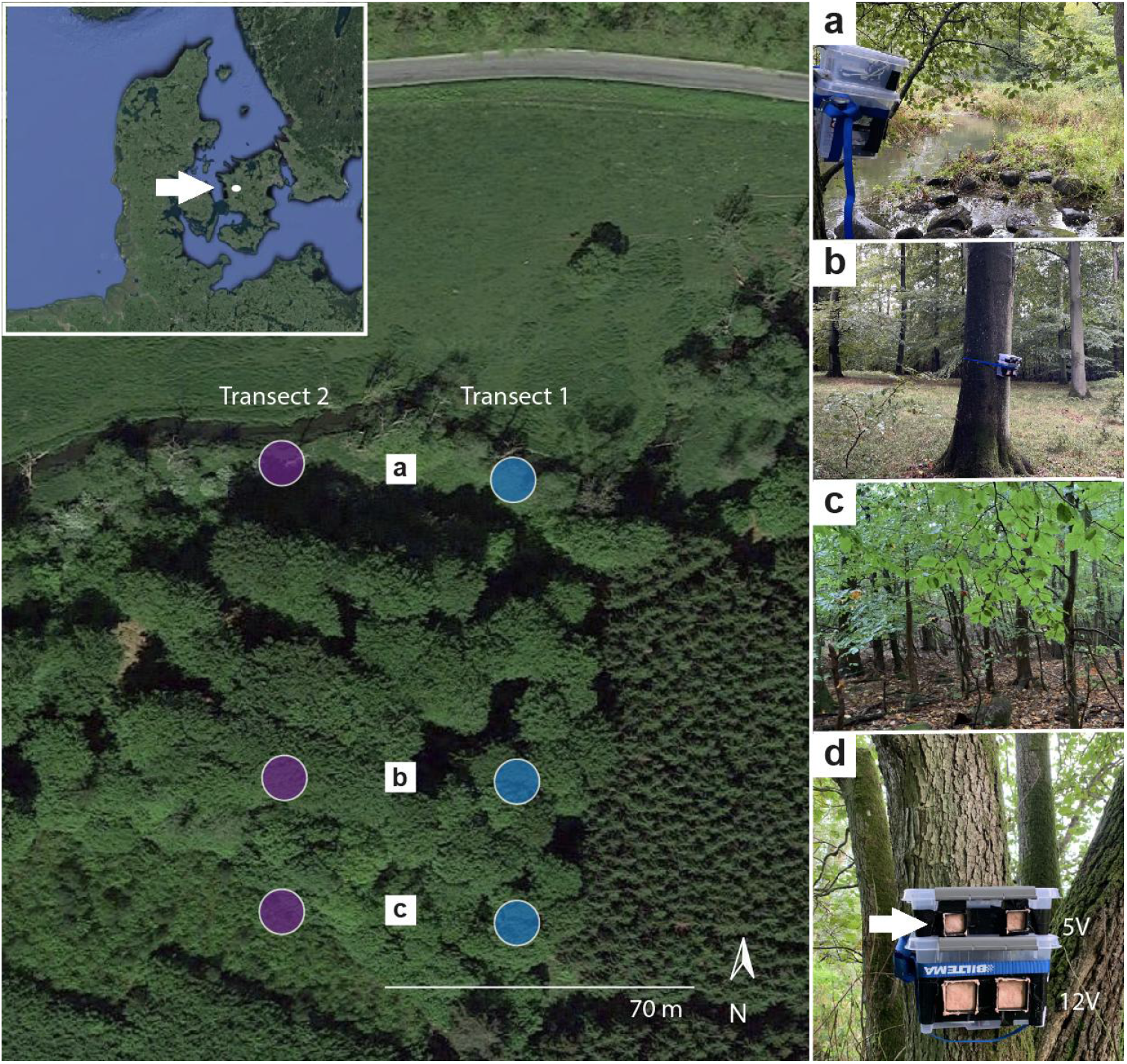
The study site in Åmosen Nature Park, Denmark, for collection of airborne environmental DNA. Samples were collected in two transects (purple and blue circles). Each transect consisted of three sampling sites spanning three microhabitat types, yielding a total of six sampling sites. The two northernmost sampling sites were located at the forest edge close to the stream (a). The next two sampling sites were located in semi-open deciduous forest (b) while the two southernmost sampling points were located in more closed deciduous forest (c). At each of the six sampling sites, two plastic boxes were fitted on a tree (d); one containing two low air flow samplers (with a 5 V fan) and one containing two high air flow samplers (with a 12 V fan). For each sampling event, a filter was fitted on each sampler (arrow). For each sampler type, simultaneous sampling yielded paired field replicates. Six 12-hour sampling events were carried out with the 24 air samplers, yielding 144 air samples.

### Vertebrate taxa detected through airborne eDNA

We used metabarcoding and high-throughput sequencing to detect vertebrate DNA within the 143 air samples. Specifically, we used two primer sets, mammal 16S ^28^ and vertebrate 12S ^29^. This resulted in the detection of 64 non-human vertebrate taxa, representing 57 wild and 7 domestic species. Overall, these comprised 4 taxonomic classes; birds (Aves), mammals (Mammalia), amphibians (Amphibia), and ray-finned fishes (Actinopterygii) spanning 17 orders and 35 families (Fig. 2). Detections included taxa known to naturally occur in Åmosen Nature Park, four taxa not previously known to occur in the area, but occurring in other areas in Denmark (stone loach (*Barbatula barbatula*), the scoter (*Melanitta* sp.), common vole (*Microtus arvalis*), and a species invasive to other European countries (Eastern grey squirrel, *Sciurus carolinensis*)). Further, we detected species exotic to the area, namely peafowl (*Pavo* sp.), budgerigar (*Melopsittacus undulatus*), cockatiel (*Nymphicus hollandicus*) and grey parrot (*Psittacus erithacus*). These are all species that are kept as pets and are known to escape into the wild in Denmark. Lastly, we detected several domestic animals, including chicken (*Gallus gallus*), turkey (*Meleagris gallopavo*), pig (*Sus scrofa*), cow (*Bos taurus*), dog (*Canis lupus*), horse (*Equus* sp) and sheep (*Ovis aries*). All the detected wild bird species were species known to occur in Denmark during the time of sampling, even accounting for migratory bird species ^30^.

**Figure 2.**
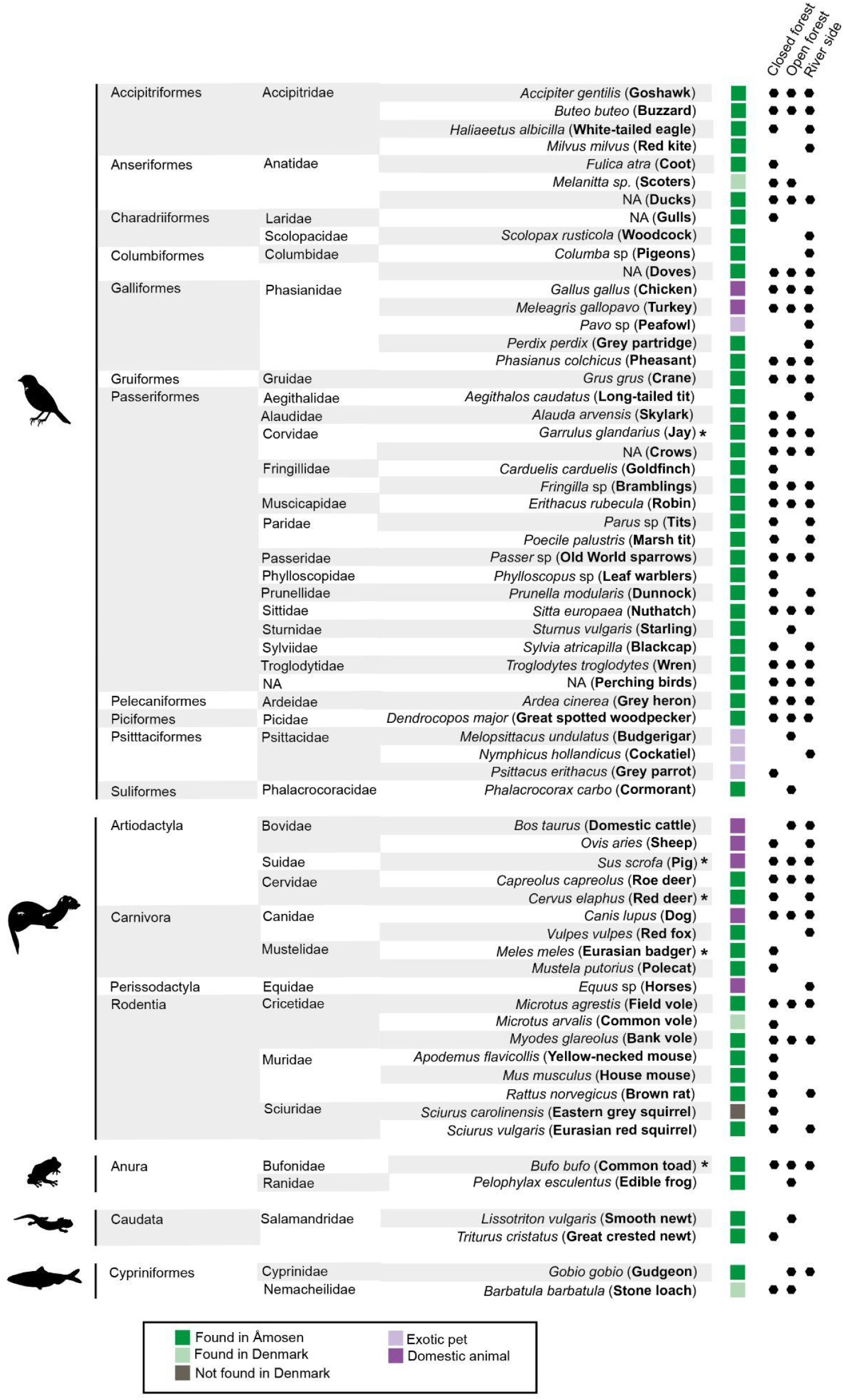
The 64 vertebrate taxa detected using metabarcoding of 143 airborne eDNA samples collected in Åmosen Nature Park, Denmark. Taxonomic order and family are listed for each taxon. Common names are listed in bold. Human detections are not included. Each taxon is characterised as belonging to one of five categories: found in Åmosen (dark green), not found in Åmosen but found in Denmark (light green), not found in Denmark (grey), exotic pet (light purple) and domestic animal (dark purple). Black dots mark which of the three micro-habitats (closed forest, open forest, and river side) each taxon was detected in. A star (*) marks that the taxon was detected with both the 16S and 12S primer sets. Common names were obtained from https://www.nhm.ac.uk/our-science/data/uk-species/index

The two metabarcoding primer sets gave complementary results with only red deer (*Cervus elaphus*), Eurasian badger (*Meles meles*), Eurasian jay (*Garrulus glandarius*), common toad (*Bufo bufo*) and pig (*S. scrofa*) overlapping between the 12S and 16S data (Fig. 2). When excluding detections of humans and domestic animals in the combined 16S and 12S dataset, 129 of the 143 samples (90.2%) yielded detections of vertebrates with 1-12 taxa detected per sample. Specifically, 31 samples had 1-3 taxa detected, 87 had 4-8 taxa and 11 samples had 9-12. The most detected taxa were birds (Aves) (Fig. 3), and the most frequently detected species was common pheasant (*Phasianus colchicus*). Despite the broad taxonomic range, sampling efficiency curves indicated that the 143 air samples collected over three days and nights did not capture the entire richness of the area’s vertebrate species. Specifically, there was no species saturation for either of the three days of sampling (merging data from day and night collections), nor when combining detections from all samples (Fig. 4).

**Figure 3.**
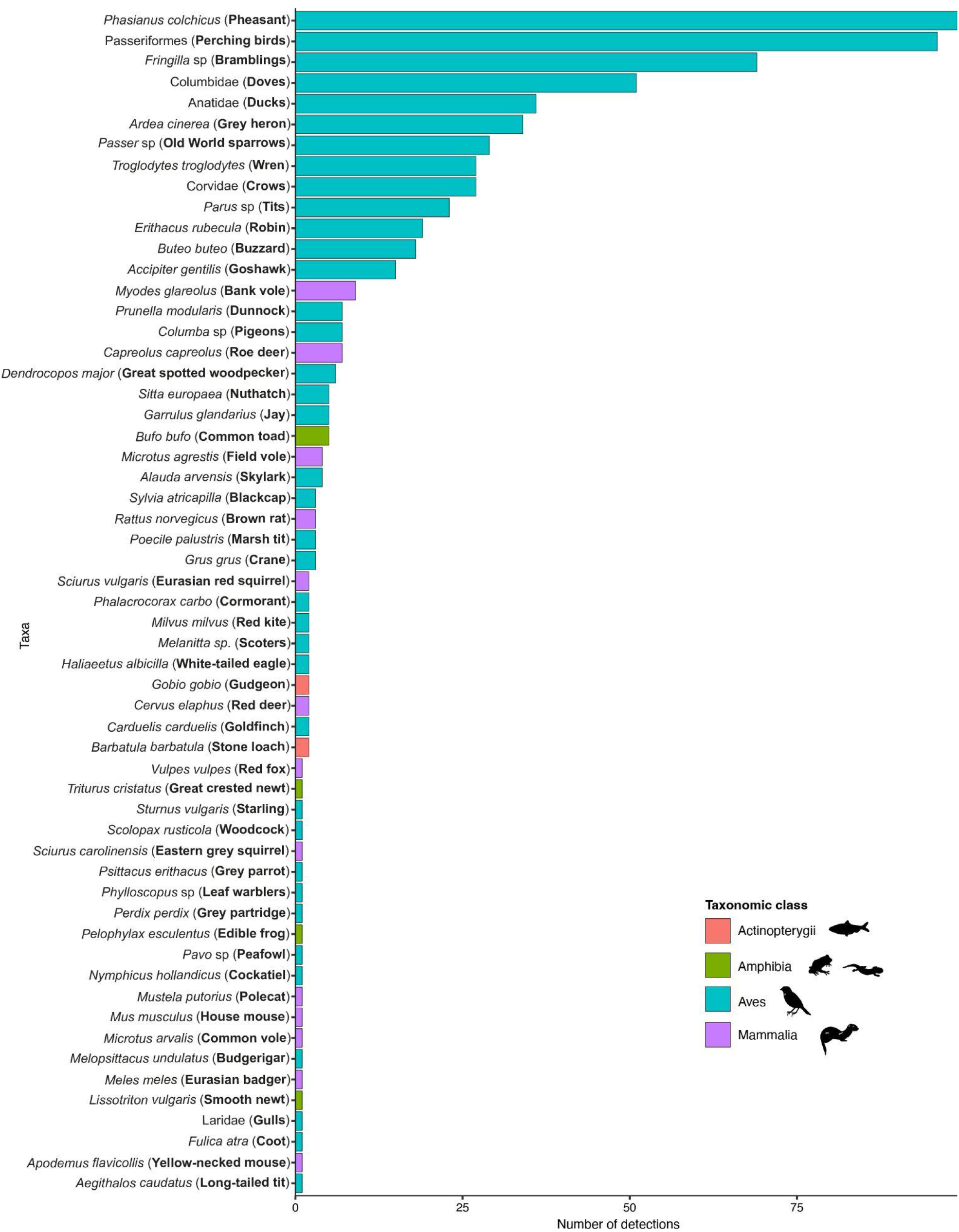
Number detections of each vertebrate taxa. Taxa were detected through metabarcoding of 143 airborne eDNA samples collected in Åmosen Nature Park, Denmark. Domestic animals and human detections are not included. Taxa are ranked by the number of samples in which they were detected. The colour of each bar indicates taxonomic class, i.e. amphibians, birds, mammals and ray-finned fish.

**Figure 4.**
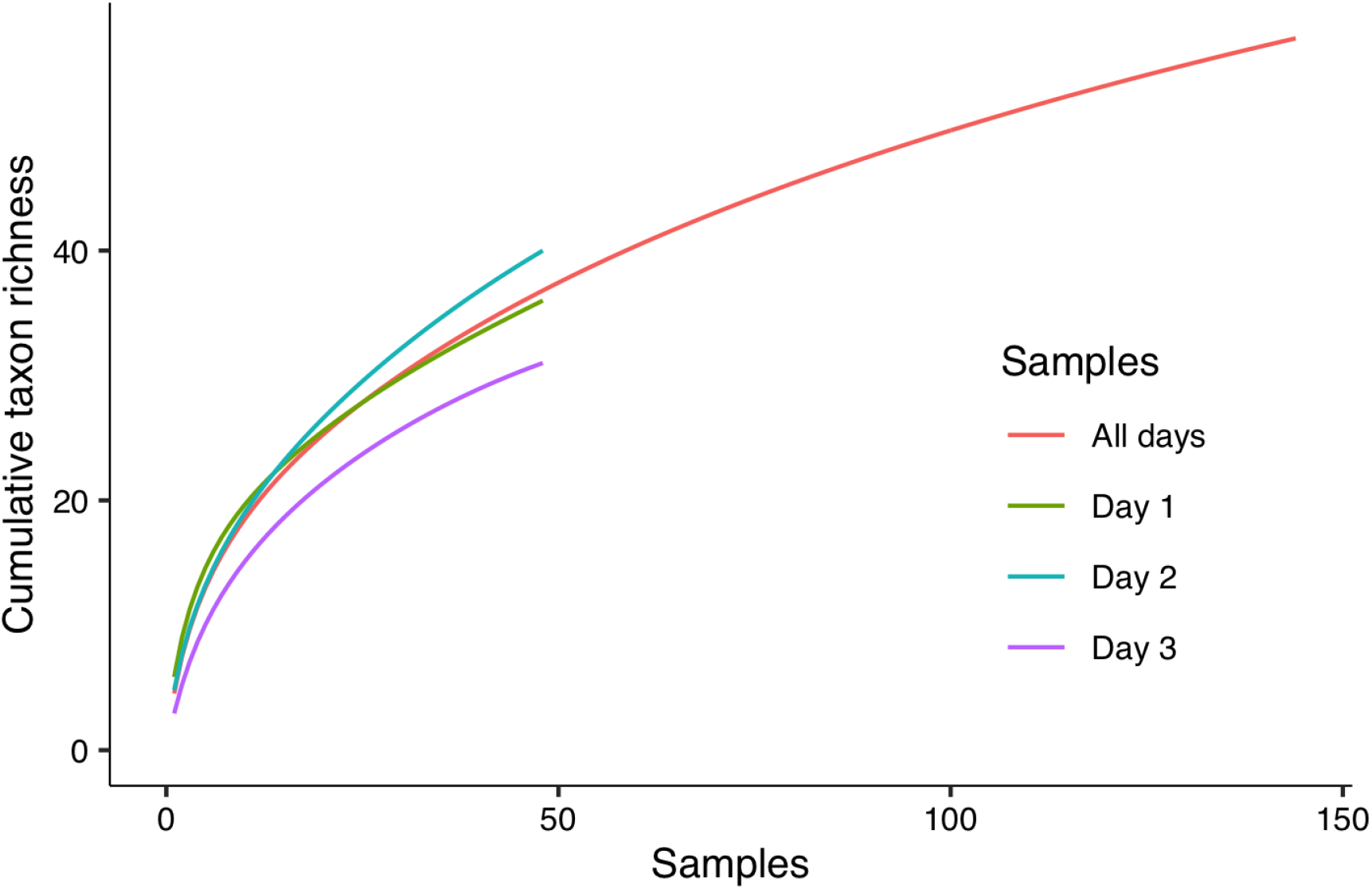
Sampling efficiency curves for vertebrate detections using metabarcoding of 143 airborne eDNA samples collected in Åmosen Nature Park, Denmark. Accumulated taxon richness is shown for each of the three sampling days (day and night sampling events combined) and for all 143 samples combined.

### Impact of field replicates and air volume

To assess the distribution of airborne eDNA and the need to include field replicates in airborne eDNA surveys in nature, we tested whether field replicates obtained using two air samplers at the same site differed in the number of taxa detected. We excluded human and domestic animals from these analyses, as our focus was on the detection of wildlife. We found no systematic difference in the number of taxa detected between paired field replicates (Wilcoxon paired test: *p* = 0.262 for the low air flow sampler and *p* = 0.7094 for the high air flow sampler). In agreement with this, the number of detected taxa were moderately correlated between field replicate pairs (*r* = 0.55 for low air flow sampler and *r* = 0.47 for high air flow sampler, both *p*-values < 0.005) (Fig. 5). To explore whether replicates within paired field replicate samples detected the same vertebrate community, we compared compositional similarities of paired field replicates for samples that had at least two taxa detections. The proportion of shared taxa (Jaccard similarity) ranged from 0 to 0.57 (median 0.30) for the low air flow sampler and from 0 to 0.75 (median 0.43) for the high air flow sampler. To explore whether field replicates were more similar to each other than to other simultaneously collected samples, we compared compositional similarities of all non-replicate samples from the same sampling event. For all non-paired samples the proportion ranged from 0 to 0.67 (median 0.27) for the low air flow sampler and 0 to 0.83 (median 0.29) for the high air flow sampler. The similarity of field replicates was significantly higher than for non-paired samples for the high air flow sampler (Wilcoxon, *p* = 0.008) and insignificantly for the low air flow sampler (Wilcoxon *p* = 0.21) (Supplementary Fig. 1a). An NMDS ordination (Supplementary Fig. 1b) using Jaccard dissimilarities (1-similarity) was used to visualise the differences in vertebrate community composition between paired field replicates for the low air flow and high air flow samplers. Only samples with two or more detections were included. This plot supports a slightly higher overall similarity between the paired field replicates, especially for the high air flow sampler.

**Figure 5.**
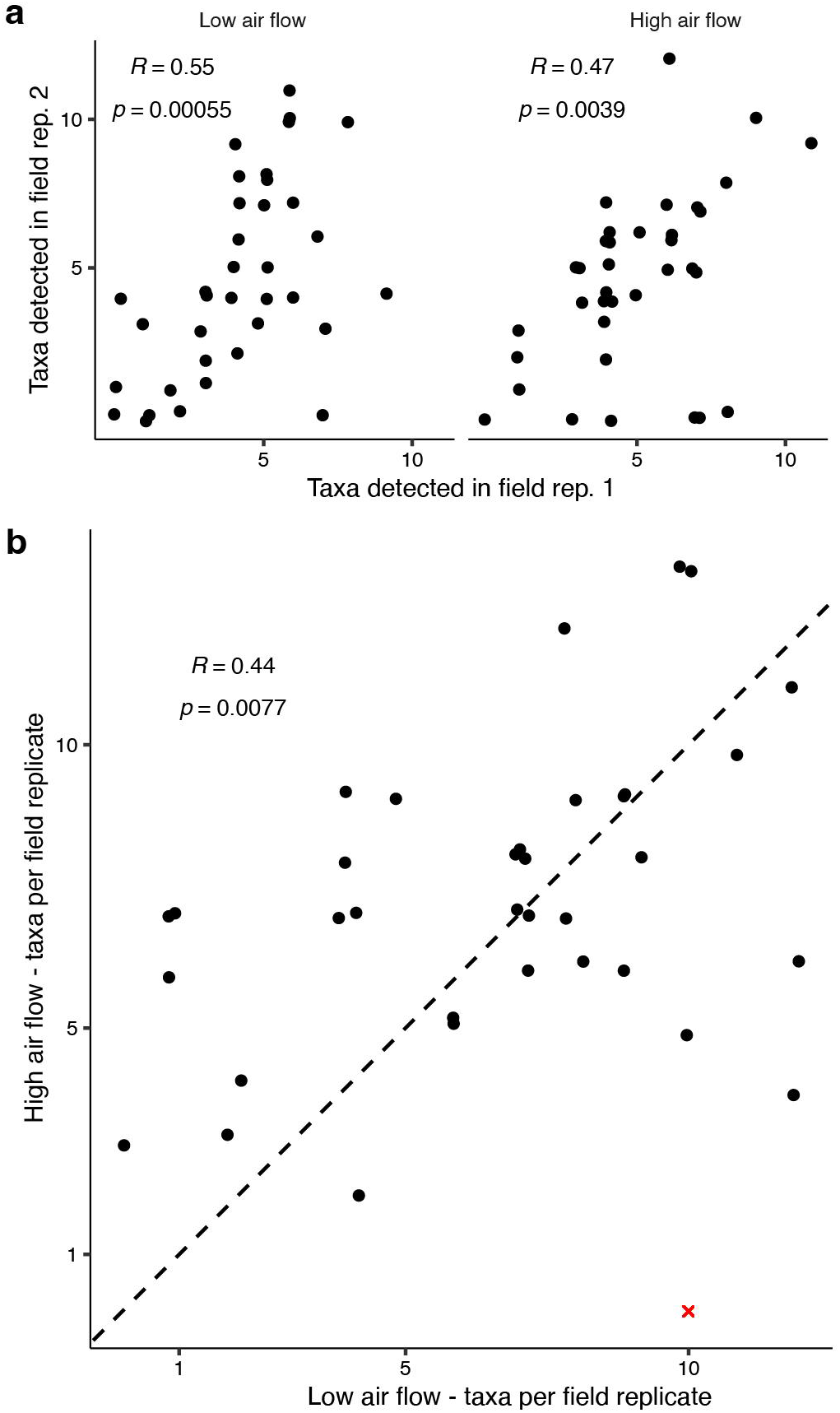
Comparisons of number of vertebrate detections between field replicates within and between low and high air flow samplers. a) Comparisons of the number of vertebrate detections in paired field replicates collected with low air flow (5 V) and high air flow (12 V) air eDNA samplers. b) Comparisons of the number of taxa detected with the two types of air samplers (low and high air flow). For each sampling site in each sampling event, data from the two paired field replicates (i.e. two high and two low flow) are combined. Detections of humans and domestic animals are excluded. The dashed line indicates the *y*=*x* line, where all low/high pairs would be in perfect correlation, i.e. have the same number of taxa detected. One outlier (marked with a red x) was removed before calculating the correlation coefficient.

We then assessed the impact of sampled air volume. Over 12 hours the low air flow sampler, operating at a flow rate of 1.1 m^3^/hr, sampled 13.2 m^3^ of air. The high air flow sampler at a flow rate of 3.5 m^3^/hr sampled 42.0 m^3^, over three times the air volume. The two air samplers had a comparable number of taxa detected per sample ranging from 0 to 11 for the low air flow sampler and from 0 to 12 for the high air flow sampler (Supplementary Fig. 2a). For both samplers, the most frequently detected number of taxa per sample was four (Supplementary Fig. 2). To further explore potential differences between the low and high air flow samplers, we combined the taxa detected by the paired field replicates. We then compared the detections for each of the six sampling sites during each of the six sampling events. Between merged field replicates collected with the low and the high air flow samplers, the number of taxa detected were significantly, although moderately, correlated (Pearson correlation test, *r* = 0.44, *p* = 0.0077, with one outlier removed) (Fig. 5b). This indicates that neither sampler is more efficient with regards to the number of taxa detected, at least for the current setup, and that successful sampling may depend more on position and air currents than on the volume of air sampled or the duration of the sampling. We further explored whether the detected vertebrate community differed between the low and high air flow samplers using Jaccard dissimilarities plotted in a NMDS ordination (Supplementary Fig. 2b). This shows that the detected community differs between the two air samplers.

### Spatial and temporal distribution of vertebrate airborne eDNA

We explored the spatial and temporal distribution of vertebrate airborne eDNA. First, for each of the six sites and six sampling events, we combined the vertebrate detections from the two field replicates collected with low air flow samplers and the two field replicates collected with the high air flow samplers, using the number of detections (0-4) of each taxon as an abundance score. This resulted in 36 compound samples. To visualise spatial and temporal dissimilarities in vertebrate composition, we calculated pairwise Bray-Curtis dissimilarities between these 36 samples and used NMDS ordinations and PERMANOVA to assess potential groupings across transect, sampling site, microhabitat, sampling event and sampling time (day vs. night) (Supplementary Fig. 3). To test for homogeneity of group dispersion, we carried out a beta dispersion test. This showed homogeneous variances when grouping the data by transect (*p* = 0.78), sampling site (*p* = 0.51), microhabitat (*p*=0.85), and sampling time (*p* = 0.29), but not when grouping by sampling event (*p* < 0.0001). The PERMANOVA analysis indicated that sampling event was the only factor with a significant effect (*p* = 0.001) on position in multidimensional space, but sampling events also showed inhomogeneous beta dispersion. Thus, we cannot say with certainty that sampling event had a significant effect on position – i.e. a significant compositional differences between sampling events – when looking at the 36 merged samples. These results indicate that, for this study setup and in this study system, vertebrate airborne eDNA was randomly distributed spatially, but may have had some temporal pattern. To further visualise the temporal pattern we combined all samples from each of the six sampling events and did a community composition analysis using the same approach as above (i.e. Bray-Curtis dissimilarity using number of detections as abundance variable) followed by a NMDS ordination (Fig. 6). This plot indicates a systematic directional temporal trend along the *x*-axis with sampling event (except for sampling event 1), indicating a systemic turnover or shift in the detected vertebrate community over time, but no apparent effect of day vs. night sampling.

**Figure 6.**
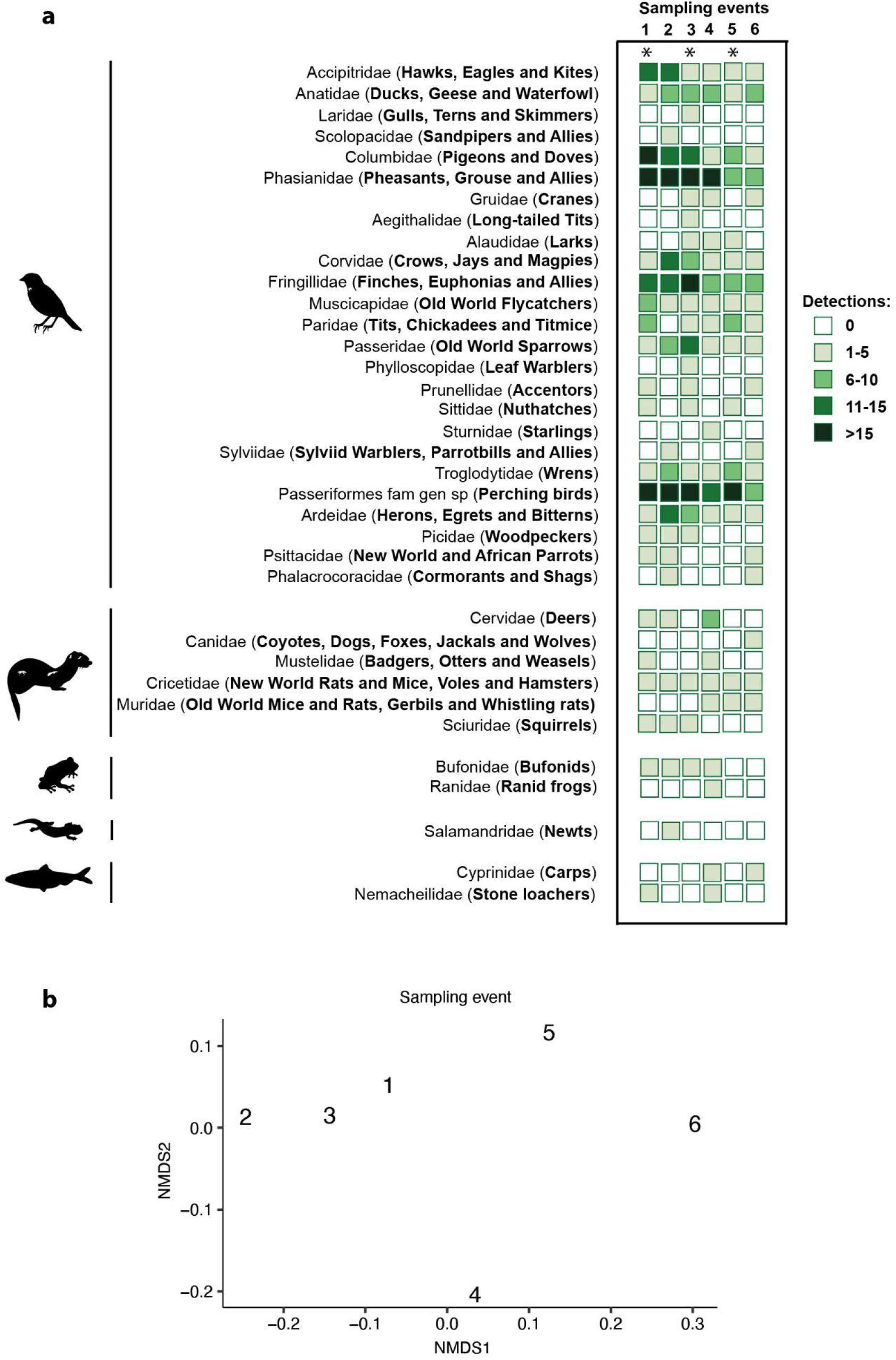
Temporal trends in vertebrate detections. (a) Taxonomic vertebrate families detected through metabarcoding of 143 airborne eDNA samples collected in Åmosen Nature Park, Denmark, in six consecutive 12-hour sampling events in the boreal autumn. The number of detections of each taxon at each of the six sampling events is shown using a colour gradient. (b) Community compositional differences of sampling events. NMDS ordination of Bray-Curtis dissimilarities of communities detected at each of the six sampling events. Number of detections was used as abundance in dissimilarity calculations. Domestic animals and humans are not included. A star (*) marks sampling events during day time. English common names taken from https://birdsoftheworld.org/ and https://animaldiversity.org/

When exploring the trajectory and origin of the sampled air, 48 hour back trajectories (Supplementary Fig. 4) showed that most of the air at the sampling site arrived from the west, and originated in the free troposphere, where we expect DNA loading will be extremely low due to oxidation, photolysis and deposition. However some of the air does arrive from the boundary layer (Supplementary Fig. 4c). As air spent the 12 hours before sampling in the planetary boundary layer, it therefore may contain aerosol DNA from the Jutland peninsula. Similarly, the air arriving from south-east (Supplementary Fig. 4f). Nonetheless, based on the data from Lynggaard ^26^, distance from source is an important variable if for no other reason than simple dilution.

To explore changes in the air particle concentration in the study site, we measured particulate matter PM10 i.e. total particle mass with an aerodynamic diameter less than 10 μm. The concentration of particles changed through time with an average of 19.78 μg/m^3^ between the first two sampling events, and decreasing up to an average of 4.72 μg/m^3^ between the fourth and fifth sampling events. After this, the concentration increased to 16.30 μg/m^3^ between the fifth and the sixth sampling (Supplementary Figure 5). Changes in the particle concentration corresponds to changes in the meteorological conditions (Supplementary Figure 6), with an increase of precipitation and wind speed and a decrease in the atmospheric pressure around the fourth sampling event.

## Discussion

We demonstrate the use of airborne eDNA for detection of vertebrate communities in nature. Through metabarcoding of airborne eDNA collected in Åmosen Nature Park, Denmark, we detected 64 bird, non-human mammal, fish and amphibian taxa spanning wild and domestic animals and exotic pets. Forty-eight of these were terrestrial vertebrate taxa naturally occurring in Denmark, i.e. wildlife (Fig. 2). Thereby, in just three days of sampling in a limited area, we detected around 20% of the approximately 210 terrestrial vertebrate species that have been registered in the greater Åmosen area through years of compiling visual observational data from various sources ^compiled by 31^. Airborne eDNA detections were randomly spatially distributed between sites, highlighting the spatial movement of airborne eDNA and the sensitivity of detecting trace DNA. A temporal pattern in the detected vertebrate communities indicated that changes in weather conditions can influence detections. The evidence from our study supports airborne eDNA as a new, and so far untapped, substrate with which to monitor a wide range of terrestrial vertebrates with limited sampling effort.

### Vertebrate taxa detected using airborne eDNA

While air sampling did not catalogue the total vertebrate richness of the area (Fig. 4), the three days of sampling in the rather limited study site provided a robust species list with identifications of 49 vertebrates known to occur in Åmosen, including one aquatic vertebrate (Fig. 2). These represent around 32.3% of the mammal, 20.1% of the bird, 57.1% of the amphibian and 4.2% of the fish species that have been observed in the area ^compiled by 31^. Importantly, OTUs that could only be identified at a higher taxonomic level were collapsed and collectively assigned to the respective genus, family or order level. This produced ten bird taxa assignments, which could represent several species (Fig. 2, 3).

The taxa we detected with airborne eDNA were ecologically, behaviourally and morphologically diverse, including carnivores, insectivores, omnivores, piscivores, and herbivores, diurnal and nocturnal animals, ground-dwelling and arboreal, volant and aquatic birds, as well as animals with fur, feathers and naked skin and spanning different body sizes. We even detected two fish species, which can be assumed to originate from foraging events or bioaerosols from faecal matter. Such diverse taxa were also identified through airborne eDNA in an urban zoo environment ^26^. However, as animals in that study were relatively numerous and confined to limited areas, those results are not directly comparable to the results we obtained in nature. For instance, in the current study, bird taxa were by far the most detected (Fig. 3), whereas more mammals than bird taxa were detected in an urban zoo ^26^. These differences may be due to the placement of the air samplers and/or the biomass and species-richness of birds and mammals in each setting, as well as features facilitating or hindering the spread of bioaerosols, such as behavioural confinement in cages, accumulation of DNA in soil and dust, bioaerosol transport by humans (e.g. zoo-keepers and visitors), and the presence of buildings or vegetation.

Existing methods for monitoring terrestrial vertebrates typically rely on observational data such as direct visual or auditory detections or camera trap data, which can make it challenging to detect species that are rare or sporadic, shy or nocturnal. Airborne eDNA enabled us to identify species in all these categories, including the elusive woodcock (*Scolopax rusticola*), the nocturnal Eurasian badger (*Meles meles*) and the sporadically occurring long-tailed tit (*Aegithalos caudatus*). We further detected a sea duck scoter genus of which two species (*Melanitta fusca* and *Melanitta nigra*) are common in Denmark, but are mainly coastal and have not previously been observed in Åmosen Nature Park ^31^. However, both species are known to migrate over land during night at the time of sampling (autumn) ^32–34^, which may have facilitated the detection in airborne eDNA.

Any method for monitoring terrestrial vertebrates has its advantages and disadvantages. For example, while direct visual or auditory detections are instant, they may require years of taxonomic training for each taxonomic group. Likewise, camera trapping also requires taxonomic expertise as well as substantial time for obtaining and processing of images. eDNA-based approaches may allow for relatively rapid sampling, but often require weeks or months of lab work, and species identifications can be limited by primer biases and DNA reference databases. Moreover, vertebrate DNA obtained from coprophagous invertebrates, such as flies, are biassed towards detection of small bodied animals ^20–23^ and only freshwater eDNA ^35^ has been found applicable for bird detection. Ultimately, as we detected a broad and diverse range of vertebrates, including birds, our results demonstrate that airborne eDNA can complement - and perhaps even replace - some existing biomonitoring methods.

### Effect of air sampling effort, timing and placement

As mentioned, the high air flow sampler collected particulate matter from 42.0 m^3^ of air during the 12 hour interval, over three times more than the 13.2 m^3^ collected by the low air flow sampler. However we did not find significant differences in the number of taxa detected by the two sampler types, nor between the two field replicates of each air sampler (Fig. 5). Nevertheless, the use of higher airflow provided a higher proportion of shared taxa between the two replicates (Supplementary Fig. 1b). This, together with the fact that no pair of field replicates detected the exact same community, indicates a patchy and scarce distribution of vertebrate airborne eDNA. It further emphasises the need for incorporation of field replicates, as is recommended for other eDNA substrate types ^36^. Studies of airborne eDNA monitoring of insects have not incorporated and contrasted field replicates ^37,38^, however in studies on marine eDNA, a similar patchy distribution of DNA has been observed with paired field replicates not detecting the exact same community ^39,40^. In the present study, we found no differences in the vertebrate composition between microhabitats, nor sampling sites. This may be due to the close proximity of sampling sites and microhabitats. Further, we did not detect any diurnal patterns in vertebrate detections, which could indicate that airborne eDNA persists longer in the environment than our 12-hour collections. In contrast to this, aquatic eDNA have reflected day-to-night differences in fish communities ^41^. However, not only is water quite a different medium than air, the sampling is also instantaneous, making it possible to observe these daily fluctuations. While sampling in consecutive 12-hour sampling events did not allow community changes to be observed in the present study, we did detect a temporal pattern in our data across the three days of sampling with detections of several taxa (e.g. pheasants and finches) decreasing on the latter sampling events (Fig. 6). While this could have been caused by differences in animal behaviour and human disturbance during air sampling, it could also have been caused by changes in weather conditions, for instance an increase in precipitation (Supplementary Fig. 6). Humidity can make airborne particles swell up and coagulate, increasing deposition of bioaerosols to the surface ^42^. Further, precipitation such as fog and falling rain can cause wet scavenging of particles, washing material out of the atmosphere. Indeed, studies have shown that rainfall can influence detection of airborne bacteria ^43^ and fungi spores ^44^ in air. Our findings support the notion that changes in the meteorological conditions affect the concentration of air particles (PM10) and therefore the vertebrate community that is detected.

The movement of air over long distances is also important to note as detected vertebrate DNA might have a different origin than the study site. This is because the lifetime of eDNA in the atmospheric environment is the result of complex dynamics including deposition with rainfall and directly to the surface, combined with denaturing due to oxidation and photolysis. In addition, air will take complex trajectories before arriving at the sampling site and therefore may contain DNA from a variety of environments. We do not expect the free troposphere to be a source of DNA, but air arriving from within the well-mixed planetary boundary layer will contain material emitted along its route. In this study, the trajectories show that some of the DNA detected in Åmosen may have been carried from Jylland (Supplementary Fig. 5c). Nonetheless, based on the data from Lynggaard ^26^, distance from source is an important variable if for no other reason than simple dilution. This is further emphasised by the fact that the vast majority of the detected species in the present study were observed in the study site (Fig. 2). As the logistical constraints of our study only allowed for a cursory exploration of the effect of meteorology on detections, this topic should be explored in future work.

### Limitations

The amount of template DNA was low in the filter samples, as illustrated by the need to use a relatively high PCR cycle number, the stochastic amplification in PCR replicates and the low overlap in species detected by the two primer sets. This was in sharp contrast to the robust detections of airborne vertebrate eDNA in an urban zoo environment^26^ and highlights the potential for optimisations to further strengthen detections in nature.

First, optimisation should focus on measures that can increase DNA template amounts. To achieve this, sampling time could be increased. Further, as DNA is sensitive to heat and UV ^45^, the spatial range of airborne eDNA might vary with season, meteorology and sunlight. To ensure authentic results and increase detected diversity, laboratory processing should follow strict lab guidelines, including incorporation of negative controls at several levels to identify sources of contamination. Further, several PCR replicates per sample could be included to optimise detection ^39^ errors and further verify detections ^46^. PCR cycle numbers and number of PCR amplification steps in the metabarcoding setup should be kept to a minimum ^46,47^. We saw very little overlap between detections made by the two primer sets (Fig. 2), highlighting both the stochastic nature of the amplifications and the need to employ complementary primer sets to increase species detections. This will also be favourable with regards to DNA reference database coverage, as species might not be represented for all markers. A further advantage is to facilitate verification of taxonomic identifications. In addition to the use of more primer sets, laboratory optimisations could include the use of blockers for non-target species amplified by the chosen primer set. For example, we obtained sequences arising from domestic animals such as pig, which may negatively affect our ability to detect naturally occurring animals. However, blockers might not fully suppress amplification, and despite the use of human blockers, we still generated human sequences.

Given the stochastic PCR amplifications in the current study, we could not use detections across PCR replicates to remove erroneous sequences ^26,46^. Instead, we pooled the sequences from each samples’ four PCR replicates and used a relatively high copy number cut-off to balance diversity detection with removal of erroneous sequences ^46^. This resulted in detections for which the occurrence in Denmark could be verified for all but one species, the grey squirrel (*Sciurus carolinensis*) (Fig. 2). Grey squirrel is an invasive species in Europe and has only once been detected in Denmark, but not in Zealand where the study was carried out ^48^. The species is, however, known to be sold and kept as a pet in Denmark. We only detected grey squirrel in one PCR replicate from one sample, but not in any of the negative controls. Nonetheless we report the detection of this species with caution, and use this potentially false positive to highlight the importance of incorporating measures to increase robustness of PCRs and ensure authenticity of airborne eDNA detections.

#### Perspectives

To inform and assess conservation efforts and to provide data for ecosystem and biodiversity studies, we need effective tools to gather data on species presence. In this study we demonstrate that terrestrial vertebrate communities can be monitored using airborne eDNA in nature. The potential of this is highlighted by that just three days of air sampling - followed by a few dedicated weeks of lab work - allowed for the detection of 22.9% of the terrestrial vertebrates observed in the area and further by the detected range of ecologically, behaviourally and morphologically diverse bird, mammal, amphibian and fish taxa. It is clear that airborne eDNA is a sensitive method with which to characterise vertebrates. However as highlighted in this study, its reliance on trace amounts of DNA introduces caveats and the need to take appropriate measures. In addition, as with any novel method, optimisations are needed to further strengthen the use of airborne eDNA for vertebrate monitoring. The study adds terrestrial vertebrates to the recent work demonstrating that plants ^49^ and insects ^37,38^ can be detected in natural systems using airborne eDNA. Thereby, airborne eDNA coupled with DNA metabarcoding can become a key tool to complement existing methods to achieve comprehensive biodiversity assessments, something which is in great demand ^50^. In particular it could become a key tool for ecosystems and taxa where a complete inventory is not easily obtained using other methods. With each new study exploring the use of airborne eDNA, we get closer to understanding its potential to revolutionise the way we monitor biodiversity.

## METHODS

### Airborne eDNA samplers

Airborne environmental DNA was collected using two custom-made air samplers of different sizes and air flows (voltages 5 V and 12 V). The low air flow (5 V) air sampler consisted of a 40 x 40 x 10 mm 5 V DC radial blower fan (Sunon, Inc) coupled to a 3D-printed housing used to hold the filter in place at approximately 3 cm from the intake of the blower fan. The low air flow air sampler was connected to a power bank, and had an airflow of 1.1 m^3^/hr as measured by their Clean Air Delivery Rate (CADR). The high air flow (12 V) air sampler consisted of a 60 x 60 x 25mm 12 V DC radial blower fan (Sunon, Inc) driven by 8 AA rechargeable batteries (Panasonic, 1.5 V) and coupled to a 3D-printed filter housing holding the filter in place at approximately 3 cm from the intake of the blower fan. The high air flow sampler provided an airflow of 3.5 m^3^/hr as measured by their Clean Air Delivery Rate (CADR). Both samplers were fitted with a class F7 glass-fibre particulate filter (SIMAS Filters A/S) normally used as a pocket air filter bag. Prior to sampling the F7 filters were cut to fit the housings of both air samplers, autoclaved, placed under UV light for 30 min and thereafter stored individually in sterile zip-lock bags.

At each sampling point in the field, duplicate sets of the low air flow and high air flow samplers were fitted in plastic boxes (Plast Team): one plastic box containing two 5 V samplers and one plastic box containing two 12 V samplers. To achieve this, holes were carved into the side of the plastic boxes to fit fans and housing. Further, ventilation holes were carved (Fig. 1). The power supplies were placed inside the plastic boxes. The days that were raining, an extra plastic lid was placed on top of the boxes to prevent the filters getting wet.

### Study site and sample collection

Air samples were collected in Åmosen Nature Park in western Zealand, Denmark (Fig. 1), characterised by a mixed landscape of forest, meadows, wetlands, agriculture, roads and smaller towns. The park’s unique habitats and vertebrate fauna is protected by two Natura2000 sites covering 80% of the park’s around 8,500 ha (https://naturparkaamosen.dk/). Approximately 262 vertebrates have been recorded in Åmosen Nature Park during the last decades (^compiled by 31^ from ^51,52^, the Danish Ornithology Society (DOF) and ^53^). Of the 262 observed species, some are classified as outliers due to presumed local extinction or because they had only been observed a few times in Denmark. The 234 species determined to be regularly occurring in Åmosen span approximately 34 mammal species, 164 bird species, 7 amphibian species, 5 reptile species and 24 fish species ^compiled by 31^.

Sampling took place in deciduous and pine forest in Åmosen Nature Park (Fig. 1, Supp. Table 1), located around 500 metres south of the 194 ha lake, Skarresø. Sampling took place in the boreal autumn from 28 September to 1 October 2022. Samples were collected in two parallel north-south transects placed approx 50 m apart, and extending from a stream at the forest edge and into the forest (Fig. 1). Samples were collected at three points in each of the two transects 30-70 metres apart, yielding six sites in total. The two northernmost sites were at the forest edge close to the stream, the next two sites were placed in semi-open deciduous forest, while the two southern-most sites were located in more closed deciduous forest (Fig. 1). On the other side of the stream was a cow pasture. At each of the six sites, two plastic boxes, one containing duplicate 5 V air samplers and one containing duplicate 12 V air samplers, were strapped to the trunk of a tree approx. 1.5 metres above ground (Fig. 1). At the two sites close to the stream, the samplers faced south. At the four remaining sites, they faced north.

Each round of sampling was approx. 12 hours in length and took place ca. 7am-7pm and ca. 7pm-7am, i.e. corresponding to the light and dark hours of the day (Supplementary Fig. 6). Six sampling events were carried out, yielding three during daytime and three during night time. Samplers were left unattended during sampling as the study site was on private land. The housings of the samplers were cleaned with ca. 5% bleach followed by 70% ethanol ^54^ to decontaminate them before and after each sampling. Medical gloves and masks were worn during sampling and the filters were handled using sterile tweezers. Exposed filters were stored individually in sterile plastic bags in a cooling box for up to 5 hours and thereafter at −20°C until DNA extraction. To ensure that taxonomic detections originated from airborne particles, filters which had gotten wet from rain despite the rain cover, were labelled upon collection so they could be omitted from further analyses.

To explore changes in air particle concentration in the study site, a TSI Model 3330 Optical Particle Sizer (OPS) was used during 1.5 hours between airborne eDNA sampling events. Meteorological conditions during the sampling was obtained from the Holbæk Denmark meteorological station ^55^ (Supplementary Figure 6).

A total of 144 air filtering samples were collected, representing two field replicates for both the low and high air flow air samplers placed in six sites for six sampling events. One filter (the second replicate of the low air flow sampler collected in the sixth sampling event in the closed forest in transect 1), which had been exposed to rain in the field, was omitted from further analyses, leaving 143 for analyses.

### DNA extraction

The DNA extraction from the filters principally followed the workflow described in ^26^. Despite this, we provide all details in the following text. To minimise contamination risk during DNA extraction, extractions were carried out in a specialised environmental DNA pre-PCR laboratory (described in ^26^). Many of the guidelines for the lab follow those used in ancient DNA laboratories, such as unidirectional workflow and the use of hair net, sleeves, facemask, two layers of medical gloves, dedicated footwear, and decontamination with 5% bleach. All steps of the extraction workflow were carried out in laminar flow hoods and using filter tips.

To avoid possible contamination due to the rain drops touching the outer edge of the filters and handling of the filters during their setup, approx. 1 cm of the edges of the filters was cut with sterile blades and discarded. After this, 3 mL and 1 mL of sterile PBS pH 7.4 (1X) (GIBCO, Thermo Fisher) was added to the filters used with the high air flow samplers and those from the low air flow samplers, respectively. With the exception of six samples (see Supplementary Information 1), DNA was extracted using the DNeasy Blood & Tissue Kit (QIAGEN, USA), following the manufacturer’s instructions with slight modifications. Due to their size, a total of 800 μL and 1600 μL of digest buffer (ATL and proteinase K) was added to the filters from the low air flow and the high air flow samplers, respectively. Moreover, DNA was eluted two times using 40 μL of EBT (EB buffer with 0.05% Tween-20 (VWR)), giving a total of 80 μl of eluted DNA. Inhibitors were removed from the eluted DNA using the OneStep PCR Inhibitor Removal kit (Zymo Research).

The negative controls included during the DNA extraction included a room control, used to test for contamination in the DNA extraction room by leaving a tube containing 50 mL sterile Milli-Q H_2_O open for approx. 48 h; a sterile filter that was not used during field work, which was used to test for contamination in the sterilisation of the filters; and finally, negative extraction controls added for every extraction round, to test for contamination in the reagents, giving a total of X samples. DNA extracts were stored in Eppendorf LoBind tubes at 20°C until further processing.

### DNA metabarcoding

The metabarcoding workflow principally followed Lynggaard et al. 2022. Despite this, it is described in detail in the following. Metabarcoding reactions were set up in a dedicated pre-PCR in which DNA extracts are not allowed, DNA was then added in the dedicated eDNA extraction lab before reactions were brought to a PCR room. Library preparations were carried out in a post-PCR lab. All steps of the metabarcoding workflow were carried out in laminar flow hoods and using filter tips.

Metabarcoding was conducted using two primer sets. To target mammals, a 16S rRNA mitochondrial primer set was used to amplify a ca. 95 bp marker; 16Smam1 (forward 5’-CGGTTGGGGTGACCTCGGA-) and 16Smam2 (reverse 5’-GCTGTTATCCCTAGGGTAACT-3’) ^28^. To target vertebrates, a 12S mitochondrial primer set was used to amplify a ca. 97 bp marker; 12SV05 forward 5’-TTAGATACCCCACTATGC-3’ and 12SV05 reverse 5’-TAGAACAGGCTCCTCTAG-’3 ^29^. The two metabarcoding primer sets are referred to as 16S mammal and 12S vertebrate, respectively. The so-called ‘tagged PCR approach’ was employed for metabarcoding ^47^: To the 5’ ends of forward and reverse primers of both primer sets, nucleotide tags of six nucleotides in length and with min. three nucleotide differences between were added. Further, one to two nucleotides were added to the 5’ end to increase complexity on the flowcell. DNA extracts from lion (*Panthera leo*) and polar bear (*Ursus maritimus*) were used as positive controls, as none of the species are found close to the sampling site in Åmosen.

Prior to tagged PCR, SYBR Green quantitative PCR, qPCR, was carried out on a subset of the sample DNA extracts. This was done to determine the optimal cycle number and DNA template volume to ensure in the following metabarcoding PCR amplifications. To achieve this, qPCRs were carried out on a dilution series (neat, 1:2 and 1:5) of the sample DNA extracts. Further, all negative controls were included in the qPCR to screen for contamination. For this, qPCRs were carried out on only undiluted extracts (2 μL template). For the 16S mammal primer set, the 20 μL reactions consisted of 2 μL DNA template, 0.75 U AmpliTaq Gold, 1x Gold PCR Buffer, and 2.5mM MgCl_2_ (all from Applied Biosystems); 0.6 mM each of 5’ nucleotide tagged forward and reverse primer; 0.2mM dNTP mix (Invitrogen); 0.5 mg/mL bovine serum albumin (BSA, Bio Labs); 3 mM human blocker (5’– 3’ GCGACCTCGGAGCAGAACCC–spacerC3)^56^ and 1 μL of SYBR Green/ROX solution [one part SYBR Green I nucleic acid gel stain (S7563) (Invitrogen), four parts ROX Reference Dye (12223-012) (Invitrogen) and 2000 parts high-grade DMSO]. The cycling parameters were: 95 °C for 10 min, followed by 50 cycles of 95 °C for 12 s, 59 °C for 30 s, and 70 °C for 25 s, followed by a dissociation curve. The 20 μL reaction mix was the same for the 12S vertebrate primer set, except for the use of a different human blocker (5’–3’ TACCCCACTATGCTTAGCCCTAAACCTCAACAGTTAAATC–spacerC3)^22^. The cycling parameters for the 12S vertebrate primer set were: 95 °C for 10 min, followed by 50 cycles of 94 °C for 30 s, 51 °C for 30 s, and 72 °C for 60 s, followed by a dissociation curve. The amplification plots from the qPCR indicated that for both primer sets, 2 μL undiluted DNA template would allow the highest template volume without PCR inhibition and that 42 and 45 cycles were optimal for the 16S mammal and 12S vertebrate primers, respectively. Some extraction controls incorporated during the laboratory work showed DNA amplification.

The metabarcoding PCRs were set up in 20 μL reactions as described above for the qPCRs, although omitting SYBR Green/ROX and replacing the dissociation curve with a final extension time of 72 °C for 7 min. The set-up included four tagged PCR replicates for each of the 143 DNA extracts, negative and positive controls, and for both primer sets. Different tag combinations were employed for each of the four PCR replicates from each sample.

Negative controls were included for every seven PCR reactions. Following amplification, PCR products were visualised on 2% agarose gels with GelRed against a 50 bp ladder. The filter negative control, PCR negative and most of the extraction negative appeared negative, with the exception of a dull band in some of the extraction negatives, while positive controls showed successful amplification. When visualised on the agarose gel, PCR replicates from individual DNA extracts did not consistently amplify. For the 16S mammal primer, 21 samples had all 4 PCR replicates amplify and 104 samples had 1-3 PCR reptiles amplify. For the 12S vertebrate primer, 82 samples had all 4 PCR replicates amplify and 54 samples had 1-3 PCR replicates amplify. The PCR products for the 12S vertebrate amplicons were pooled if at least one of the four PCR replicates showed a band when visualised on agarose gel. This resulted in 137 pooled and sequenced samples. For the PCR products of the 16S mammal primer, to avoid a low concentration for library preparation, PCR products were pooled only if at least two of the PCR replicates showed a positive amplification. For 16S mammal, 82 samples were pooled and sequenced.

The amplicon pools included samples, as well as all negative and positive controls. The field replicates were processed separately and therefore the pooling resulted in eight amplicon pools for each primer set: one pool for each of the four PCR replicates and for each of the two field replicates. The 16 amplicon pools were purified with MagBio HiPrep beads (LabLife) using a 1.6x bead:amplicon pool ratio and eluted in 35 μL EB buffer (QIAGEN). Library preparation of the purified amplicon pools was performed with the TagSteady protocol to avoid tag-jumping ^57^. The eight libraries were dual-indexed with indexes of 7 nucleotides in length. The libraries were purified with MagBio HiPrep beads (LabLife) with a 1.6x bead: library ratio, eluted in 30 μL EB buffer and the purified libraries were qPCR quantified using the NEBNext Library Quant Kit for Illumina (New England BioLabs). The qPCR results guided equimolar pooling of the libraries prior to sequencing on an Illumina MiSeq sequencing platform using v3 chemistry at the GeoGenetics Sequencing Core, University of Copenhagen, Denmark. 15% PhiX were included and 300 bp paired-end sequencing were carried out aiming at 30,000 paired reads for each of the four PCR replicates of each primer set, equalling an estimated 120,000 reads per sample. The libraries of the field replicate 1 were sequenced separately from the field replicate 2.

### Data analyses

Processing of sequence data principally followed ^26^. It is outlined in the following text. The sequence data from the 16s mammal and 12s vertebrate were processed separately. The sequenced data from the two field replicates were processed separately. AdapterRemoval v2.3.1 ^58^ was used to remove Illumina adapters and low quality reads and to merge paired reads. Min length was set to 100, min alignment length to 50, min quality to 28 and quality base to 33. Within each amplicon library, Begum ^59^ was used to sort sequences based on primer and 5’ nucleotide tag sequences. For primer identifications, two mismatches were allowed. Begum was further used to filter sequences across each sample’s four PCR replicates. Due to the stochasticity seen in the PCR amplification, sequences present in minimum one of a sample’s four PCR replicates and present in minimum 30 and 35 copies were retained for the 16s mammal and 12s vertebrate datasets, respectively. The datasets of the two field replicates were combined and the filtered sequences were clustered into operational taxonomic units (OTU) using SUMACLUST with a similarity score of 97% ^60^. OTU sequences were assigned taxonomy through manual ‘blastn’ searches against NCBI GenBank in which reference data from Danish vertebrates are included as part of the DNAmark project ^61^. Doing so, species-level identification was assigned if the OTU sequence had a 99-100% identity match, the highest query coverage and lowest e-value to a NCBI reference sequence.

For the 12S vertebrate data, many OTUs with less than 250 copies did not have a 100% query coverage in the blast results. Further, they matched to several taxonomic genera. Therefore, the detections with less than 250 copies were removed from further analysis as they were considered likely errors. This caused potentially true detections (i.e. sheep (*Ovis aries*), night heron (*Nycticorax nycticorax*) and golden plover (*Pluvialis apricaria*) to be removed from the data set. For OTU sequences with 100% matches to more than one species, we manually searched the geographical distribution of the species, and assigned it to the one found in the study area. For example, an OTU that matched several species of the genus *Pelophenax*, was assigned to the edible frog (*Pelophenax esculentus*) as it is the only species of *Pelophylax* that occurs in Åmosen Nature Park. However, the species has a hybrid origin from *Pelophenax ridibundus* and *Pelophenax lessonae*, and the three species cannot be distinguished with 12S sequence data ^62^. In addition, OTUs that differed only by minor length differences were merged. This was the case for e.g. *Columba* sp, *Phasianus colchicus, Capreolus capreolus* and *Vulpes vulpes*.

We verified the known distributions of the detected taxa and assessed the likelihood of their actually being detected in the study area at the sampling time by searching for the taxa individually on the Global Biodiversity Information facility (GBIF.org), arter.dk, dofbasen.dk,, a report from the Danish ornithology society ^63^, and a compilation of observation data by Holm ^31^. The migration pattern and presence in Denmark of detected bird species was verified through the Danish Bird Migration Atlas ^34^.

To reduce amplification of human DNA, human blockers were used with both primer sets. Despite this, human 12S sequences were detected in 29 samples. Of the total number of OTU sequences in these samples, human sequences comprised an average of 6.05% (range 0.02-44.48%). For the 16S mammal marker, human sequences were detected in 24 samples with an average of 8.14% (range 0.13-29.60%) of the total number of OTU sequences in these samples. Pig sequences were detected in 76 samples with the 16S mammal primers, comprising an average of 75.49% of the total number of OTU sequences in these samples (range 0.06-100%). In the 12S vertebrate data, pig sequences were detected in 10 samples with an average of 1.47% (range 0.04-3.60%) of the total OTU sequences in these samples. The detection of pig DNA could result from the multiple pig farms in the area ^64^, as well as the common use of pig manure as fertiliser on agricultural fields.

In addition, OTUs detected in the positive controls were not found in air samples. In the 16S mammal dataset, OTUs identified as human were detected both in the negative extraction controls and in some samples. In addition, pig was detected in a negative extraction control, in the room control and in most of the collected air samples. The amount of pig reads in the controls ranged from 39 to 70, but from 34 to 183,124 in the samples. In the 12S vertebrate dataset, OTUs identified as Cyprinidae and human were detected in both the negative extraction controls and samples. OTUs detected in the negative controls and those identified as domestic animals (chicken, turkey, pig, cow, dog, horse and sheep) were removed from the dataset before downstream analyses. The grey squirrel (*Sciurus carolinensis*) was detected in one PCR replicate from one sample. Grey squirrel faecal samples had previously been processed in another lab within the same overall facility where the air samples were processed, which may have caused contamination. Nevertheless, we did not detect grey squirrel in any of the negative controls.

### Statistical analyses

We counted the number of taxa detected in each sample, and the number of detections was assessed for each taxon, and congruence between paired field replicates and between sampler type was visualized with histograms, dotplots and sampling efficiency curves, and statistically assessed by Wilcoxon tests. Analyses were carried out with the functions specaccum, vegdist, betadisper, metaMDS from the r-package vegan ^65^.

Sampling efficiency curves (cumulative taxon richness as a function of the number of samples) were calculated for both the full data and per day ^66^. Compositional differences in the detected communities were assessed by calculating community (dis-)similarity measures, and visualising these with histograms and NMDS ordinations. For compositional similarity of two samples we used the Jaccard dissimilarity metric, that measures the proportion of the community that is shared by both samples (intersection over union). For the ordination analyses of single samples we used the Jaccard dissimilarity (1-similarity) and NMDS carried out with the metaMDS function of vegan (using k=2, try = 2000, trymax = 2000). We merged paired field replicates (for a total of 72 merged samples), and visualised the compositional differences between sampler types (deployed at the same time – i.e. same sampling site and event) with NMDS using Jaccard dissimilarities.

To evaluate overall temporal and spatial patterns, we merged the data from all four filters simultaneously deployed at one sampling site in each sampling event (for a total of 36 merged samples), and used the number of detections per taxon (0 to 4) in each as an abundance score. For compositional analyses of merged samples we used the Bray-Curtis dissimilarity measure. We then tested whether the community dissimilarities showed equal dispersion (beta-dispersion) across all groups in major subdivisions of the samples – transect, sampling site, sampling event and microhabitat – using the betadisper function (library vegan) testing each variable separately. We then tested for effects of location in ordination space using PERMANOVA as implemented in the adonis2 function, again using transect, sampling site, sampling event and microhabitat as potential explanatory variables, using marginal testing. Finally, we visualised the community dissimilarities with NMDS ordination as above. As sampling event seemed to be the only variable with a systematic effect on the taxa composition, we merged all detections per sampling event (for a total of 6 merged samples, each with 0-24 detections per sample). Thereafter we did a community dissimilarity analysis using NMDS ordination to visually inspect whether there was a compositional trend with time using the Bray-Curtis dissimilarity as above.

### Particle concentration measurements and air modelling

To explore changes in the concentration of airborne particulate matter with an aerodynamic diameter less than 10 μm (PM 10) in the area, air samples were taken with a TSI Model 3330 Optical Particle Sizer (OPS). The OPS was placed in the middle of the study area and left running for 1.5 hrs while the air filters were being changed at the different sampling sites. This was done each time filters were changed (i.e. between sampling events 1 and 2, 2 and 3, 3 and 4, 4 and 5, 5 and 6) and gave a total of five sampling events. Thereafter, the average concentration of all the measurements taken in each sampling event was calculated (Supplementary Fig. 5).

Atmospheric back trajectories from the air arriving at Åmosen Nature Park during the sampling events were modelled using the HYSPLIT model ^67^ and the National Centers for Environmental Prediction (NCEP) Global Data Assimilation System (GDAS) 1 degree global meteorology dataset ^68,69^.

## Acknowledgements

We thank Jacob Agerbo Rasmussen for assistance with the design and printing of 3D housings for the air samplers, Peter Rask Møller for discussions regarding choice of study area, Sarah S. T. Mak for discussions regarding laboratory protocols, Tina Brand, Pernille Selmer Olsen and Lasse Vinner (Globe Institute Molecular Biology Labs and GeoGenetics Sequencing Core) for support, infrastructure, discussions and sequencing and Tom Gilbert for comments on the original manuscript. This work was primarily supported by a VILLUM FONDEN Experiment grant awarded to K.B. and M.T.O. (00028049). Further, it was supported by a Carlsberg Foundation Young Researcher Fellowship awarded to K.B. (CF21-0411). C.L. was further supported by a research grant (41390) from VILLUM FONDEN. Animal silhouettes used in figures were obtained from the Integration and Application Network (https://ian.umces.edu/media-library/). The authors acknowledge the NOAA Air Resources Laboratory (ARL) for the provision of the HYSPLIT transport and dispersion model and READY website (https://www.ready.noaa.gov) used in this publication.

## Author contributions

Conceiving the study: C.L., M.S.J., M.T.O., K.B. Sampling permit: M.T.O. Data collection: C.L., M.T.O., K.B. Data generation: C.L. Data analysis: C.L., T.F., M.S.J. Interpretation of data: C.L., M.T.O, K.B., T.F., M.S.J. Writing the original manuscript: C.L., K.B. Revision of the manuscript: All authors. All authors approved the submitted version of the manuscript.

## Declarations of interests

M.S.J. is Chief Science Officer at AirLabs. The current study is not of direct commercial value to AirLabs. The remaining authors declare that they have no competing interests.

## Notes

### Summary of Updates

Reference to Holm et al (MSc thesis, reference number 31) replaced by reference to biorxiv preprint.

