## Supporting information for "Airborne environmental DNA captures terrestrial vertebrate diversity in nature"

### SUPPLEMENTARY INFORMATION

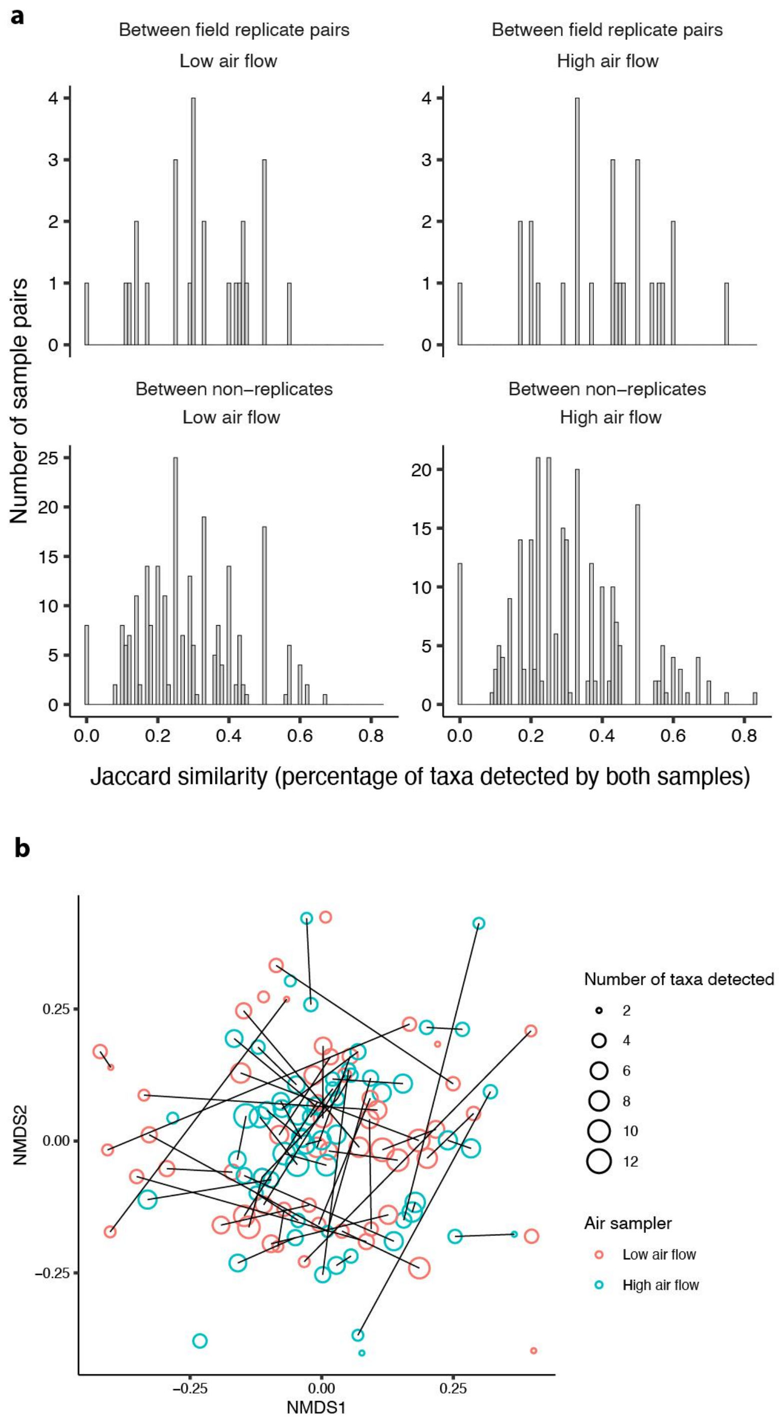


**Supplementary Figure 1**. **Pair-wise comparisons of taxa detected in airborne eDNA field replicates collected in Åmosen Nature Park, Denmark**. (a) Pairwise sample similarity. Histograms show the percentage of shared taxa detections (Jaccard similarity) between two samples from the same sampling event. Detections of domestic animals and humans are not included. Only samples with two or more taxa detections are included. The upper row shows similarities between paired field replicates, while the lower row shows similarities between non-paired samples. The left column shows pair-wise similarities for the low air flow sampler, and the right column for the high air flow sampler. The similarities of field replicates (top row) are significantly higher than values of non-replicate comparisons for the high air flow sampler (Wilcoxon $p$ = 0.008), but not for the low air flow sampler (Wilcoxon $p$ = 0.21). The similarity of high air flow field replicate pairs is significantly higher than the similarity of the corresponding low air flow replicate pairs (paired Wilcoxon $p$-value = 0.008). (b) NMDS ordination of vertebrate community using Jaccard dissimilarities between paired field replicates for the high and low air flow samplers. Detections of domestic animals and humans are not included. Only samples with two or more taxa detections are included. Field replicates are connected by lines.

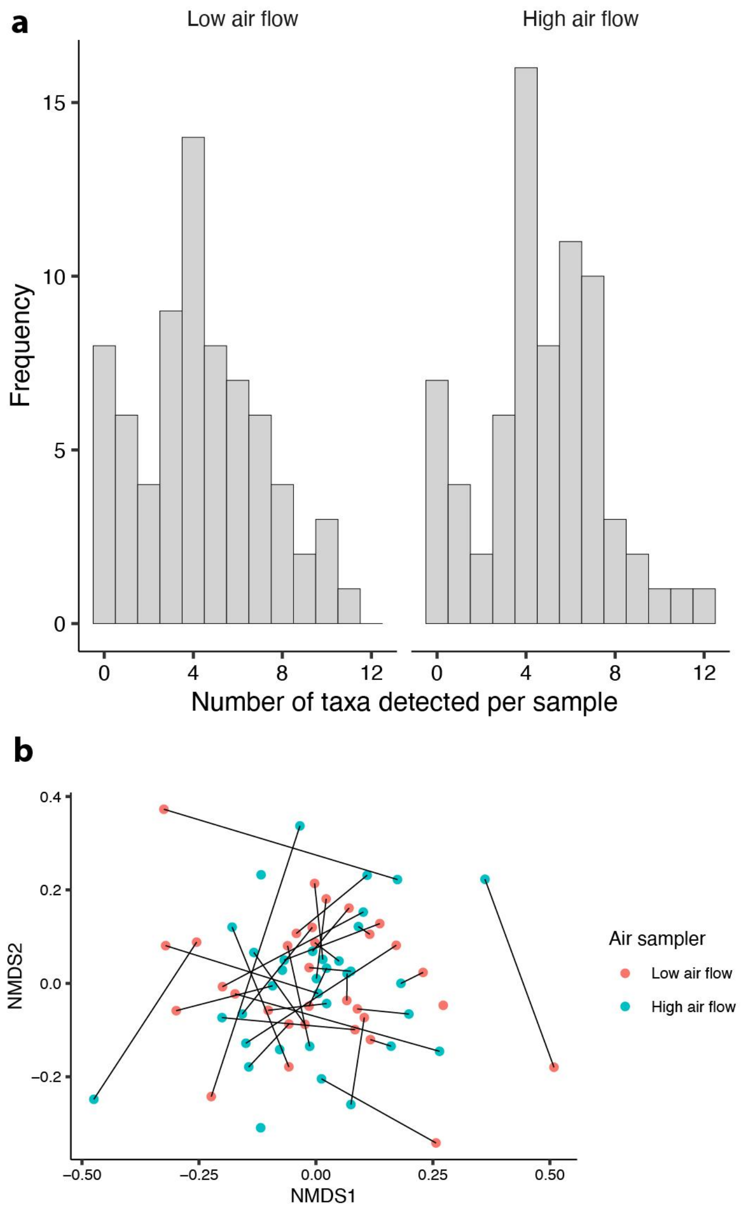

**Supplementary Figure 2. Comparison of taxa detected between the low and high air flow samplers used for airborne eDNA collection in Åmosen Nature Park, Denmark.** a) Number of vertebrate taxa detected with the two air samplers. b) NMDS ordination of Jaccard dissimilarities in the detected vertebrate community. Low and high air flow samplers from the same transect, microhabitat and sampling event are connected by a line. Data from field replicates is merged. Detections exclude humans and domestic animals.

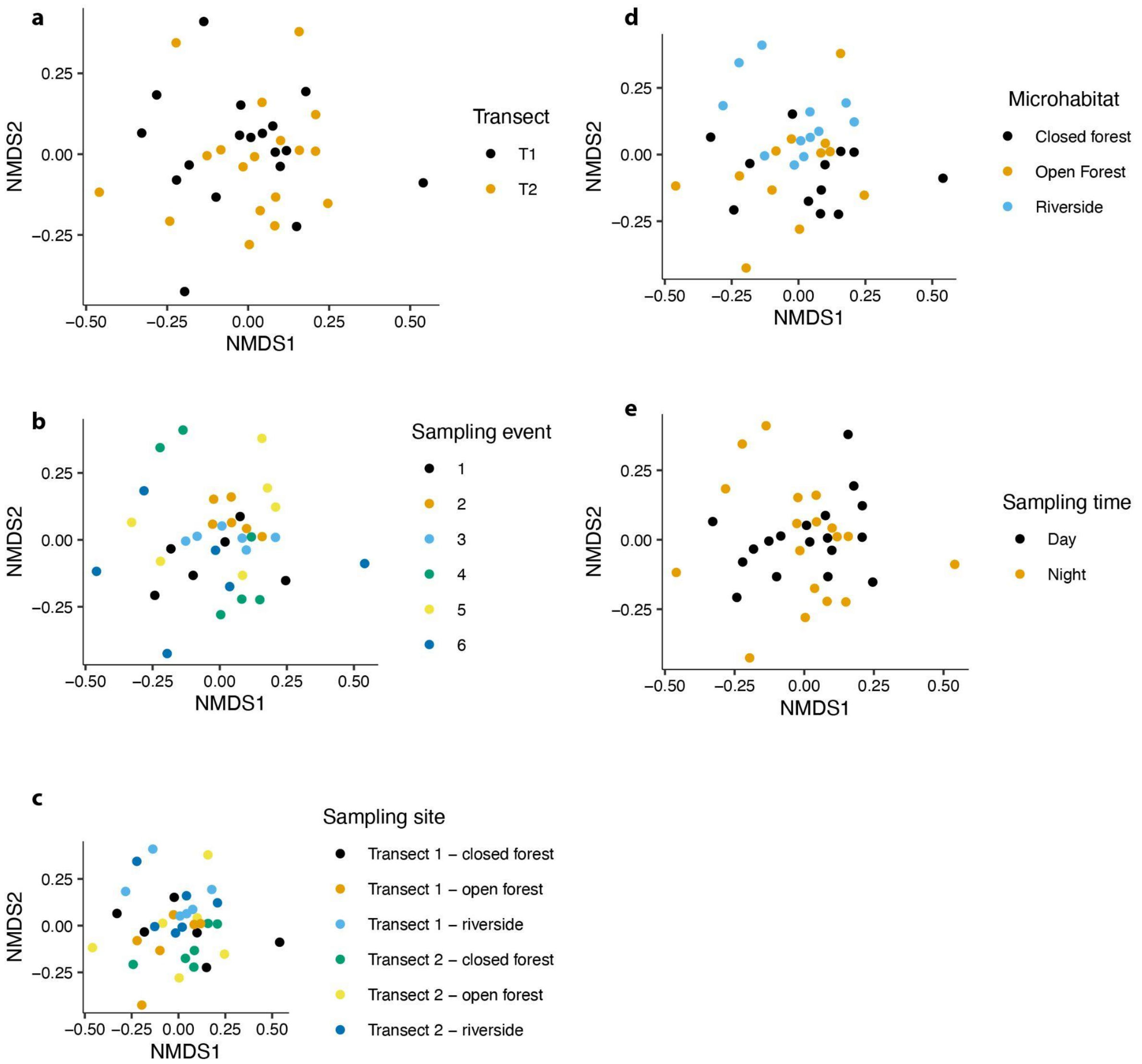


**Supplementary Figure 3. NMDS ordination of pairwise Bray-Curtis dissimilarities of vertebrate taxa compositions.** For each of the six sites and six sampling events the two low and two high air flow field replicates were combined before visualisation. a-e) show the same ordination but with colours representing different groupings of the data (a) transect, b) sampling event, c) sampling event, d) microhabitat and e) sampling time (day vs. night).

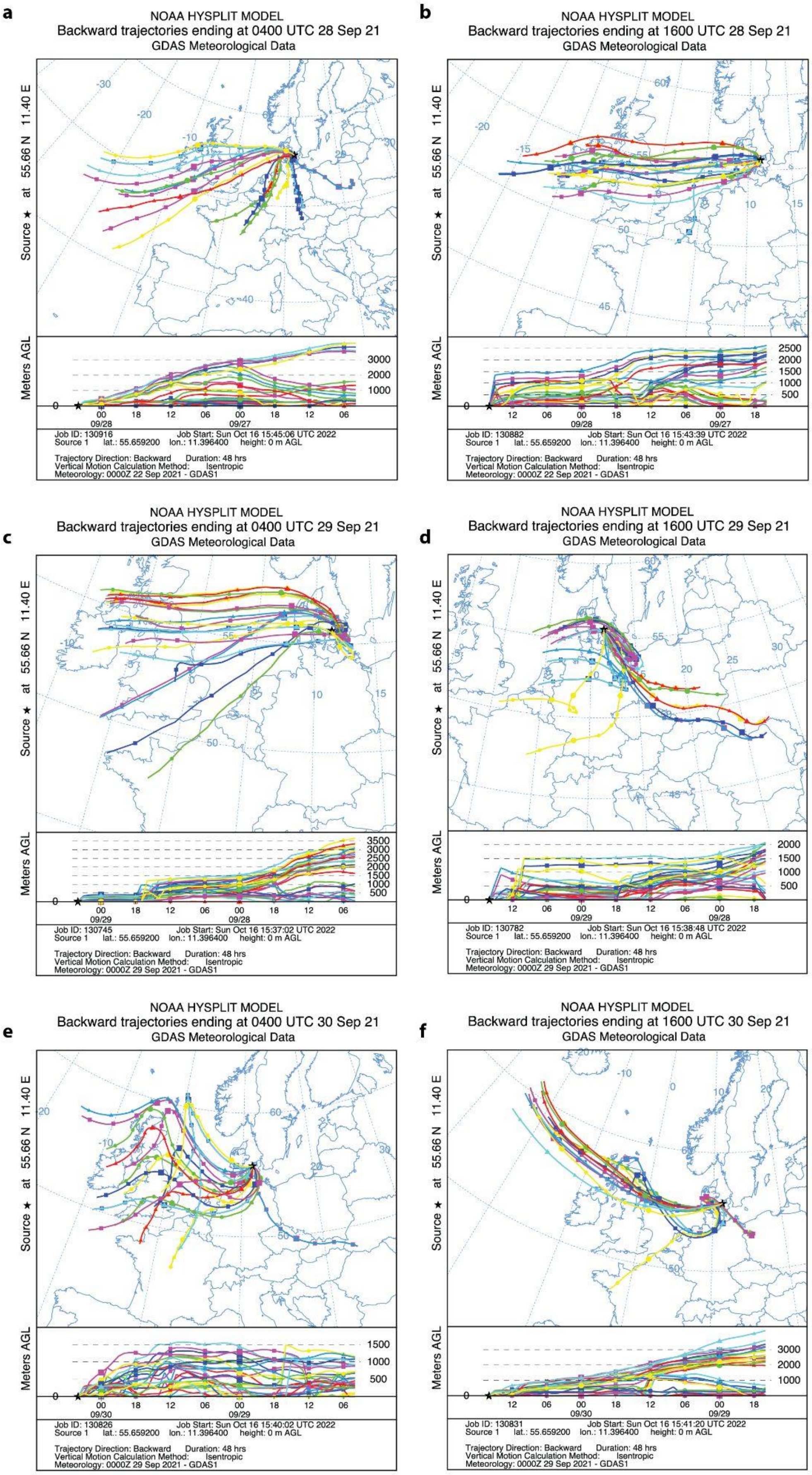
a
NOAA HYSPLIT MODEL
Backward trajectories ending at 0400 UTC 28 Sep 21
GDAS Meteorological Data
Source ★ at 55.66 N 11.40 E
Meters AGL
Job ID: 130916 Job Start: Sun Oct 16 15:45:06 UTC 2022
Source 1 lat.: 55.659200 lon.: 11.396400 height: 0 m AGL
Trajectory Direction: Backward Duration: 48 hrs
Vertical Motion Calculation Method: Isentropic
Meteorology: 0000Z 22 Sep 2021 - GDAS1
b
NOAA HYSPLIT MODEL
Backward trajectories ending at 1600 UTC 28 Sep 21
GDAS Meteorological Data
Source ★ at 55.66 N 11.40 E
Meters AGL
Job ID: 130882 Job Start: Sun Oct 16 15:43:39 UTC 2022
Source 1 lat.: 55.659200 lon.: 11.396400 height: 0 m AGL
Trajectory Direction: Backward Duration: 48 hrs
Vertical Motion Calculation Method: Isentropic
Meteorology: 0000Z 22 Sep 2021 - GDAS1
c
NOAA HYSPLIT MODEL
Backward trajectories ending at 0400 UTC 29 Sep 21
GDAS Meteorological Data
Source ★ at 55.66 N 11.40 E
Meters AGL
Job ID: 130745 Job Start: Sun Oct 16 15:37:02 UTC 2022
Source 1 lat.: 55.659200 lon.: 11.396400 height: 0 m AGL
Trajectory Direction: Backward Duration: 48 hrs
Vertical Motion Calculation Method: Isentropic
Meteorology: 0000Z 29 Sep 2021 - GDAS1
d
NOAA HYSPLIT MODEL
Backward trajectories ending at 1600 UTC 29 Sep 21
GDAS Meteorological Data
Source ★ at 55.66 N 11.40 E
Meters AGL
Job ID: 130782 Job Start: Sun Oct 16 15:38:48 UTC 2022
Source 1 lat.: 55.659200 lon.: 11.396400 height: 0 m AGL
Trajectory Direction: Backward Duration: 48 hrs
Vertical Motion Calculation Method: Isentropic
Meteorology: 0000Z 29 Sep 2021 - GDAS1
e
NOAA HYSPLIT MODEL
Backward trajectories ending at 0400 UTC 30 Sep 21
GDAS Meteorological Data
Source ★ at 55.66 N 11.40 E
Meters AGL
Job ID: 130826 Job Start: Sun Oct 16 15:40:02 UTC 2022
Source 1 lat.: 55.659200 lon.: 11.396400 height: 0 m AGL
Trajectory Direction: Backward Duration: 48 hrs
Vertical Motion Calculation Method: Isentropic
Meteorology: 0000Z 29 Sep 2021 - GDAS1
f
NOAA HYSPLIT MODEL
Backward trajectories ending at 1600 UTC 30 Sep 21
GDAS Meteorological Data
Source ★ at 55.66 N 11.40 E
Meters AGL
Job ID: 130831 Job Start: Sun Oct 16 15:41:20 UTC 2022
Source 1 lat.: 55.659200 lon.: 11.396400 height: 0 m AGL
Trajectory Direction: Backward Duration: 48 hrs
Vertical Motion Calculation Method: Isentropic
Meteorology: 0000Z 29 Sep 2021 - GDAS1

**Supplementary Figure 4. Modelled atmospheric back trajectories showing the origin of air arriving at Åmosen Nature Park during the time of airborne eDNA sampling.** Times are indicated as Coordinated Universal Time, add two hours to obtain the local time at Åmosen. 48 hour isentropic back trajectory showing origin of air arriving a 0 metres above ground level at the specified time, using the HYSPLIT model (https://www.ready.noaa.gov/HYSPLIT.php) and the National Centers for Environmental Prediction (NCEP) Global Data Assimilation System (GDAS) 1 degree global meteorology dataset [67,68].

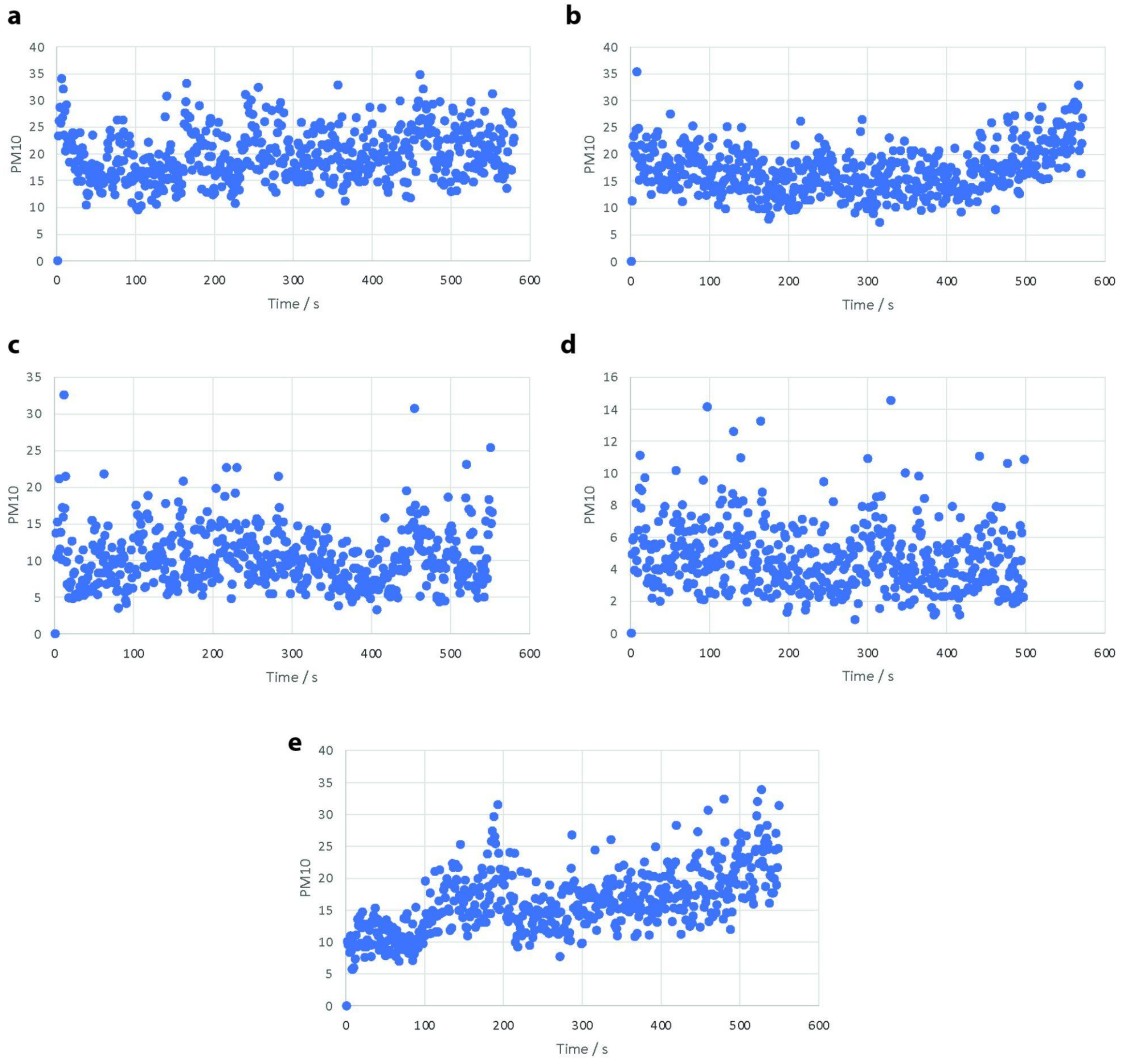


**Supplementary Figure 5. Measurements showing the concentration of PM10, i.e. total particle mass with an aerodynamic diameter less than 10 µm, during airborne eDNA collection in Åmosen Nature Park, Denmark.** Particle density assumed to be 1 g/ml. Data obtained using a TSI Model 3330 Optical Particle Sizer (OPS). (a) 2021/09/28, starting 17:32:12 CET, average PM10 19.78 µg/m$^3$. (b) 2021/09/29, starting 07:22:21 CET, average PM10 17.01 µg/m$^3$. (c) 2021/09/29, starting 17:33:11 CET, average PM10 10.29 µg/m$^3$. (d)

2021/09/30, starting 07:22:40 CET, average PM10 4.72 µg/m$^3$. (e) 2021/09/30, starting 17:30:25 CET, average PM10 16.30 µg/m$^3$.

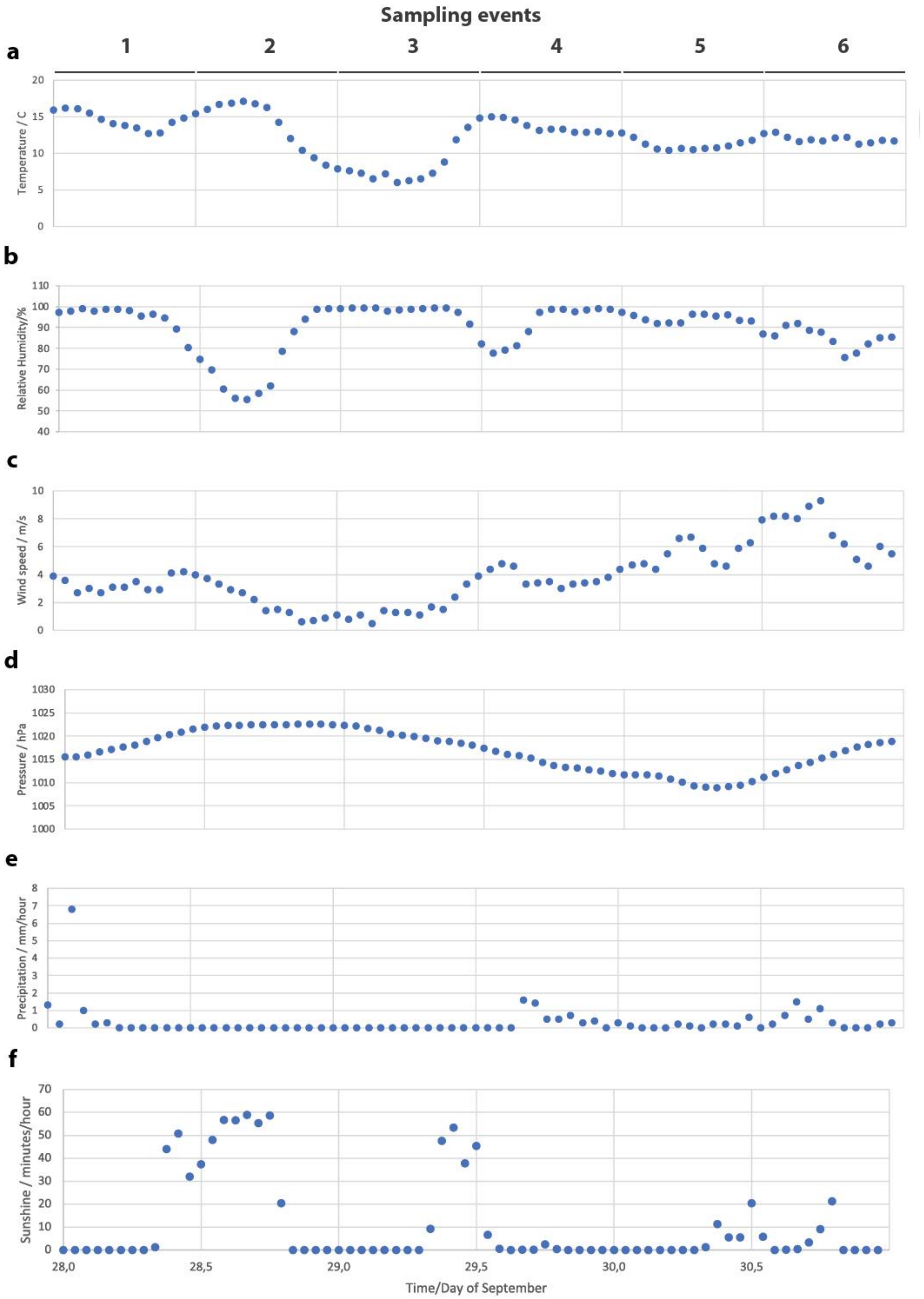

**Supplementary Figure 6. Meteorological conditions during the sampling of airborne eDNA in Åmosen Nature Park, Denmark.** Data from Holbæk Denmark meteorological station from 28, 29 and 30 September 2021. The six 12-hour sampling events are indicated as 1-6 at the top of the figure. (a) Hourly average temperature, (b) Hourly average relative humidity. (c) Hourly average wind speed. (d) Atmospheric pressure. (e) Precipitation. (f) Sunshine. Data archived at https://www.dmi.dk/vejrarkiv/, accessed 15 October 2022.

**Supplementary Table 1. GPS and sampling times for collection of airborne eDNA in Åmosen Nature Park, Denmark.** For each of the six sampling events, the GPS position for each of the six sampling sites is provided alongside information on microhabitat, date and time for placement and collection of the air filters.

| Transect | Micro-habitat | GPS position | Sampling event | Date and time of filter placement | Date and time of filter collection |
|---|---|---|---|---|---|
| T1 | Closed forest | 55°38'19"N 11°22'9"E | 1 | 28/09/21 7:37 am | 28/09/21 5:40 pm |
| | | | 2 | 28/09/21 5:53 pm | 29/09/21 7:28 am |
| | | | 3 | 29/09/21 7:40 am | 29/09/21 5:41 pm |
| | | | 4 | 29/09/21 5:52 pm | 30/09/21 7:30 am |
| | | | 5 | 30/09/21 7:40 am | 30/09/21 5:37 pm |
| | | | 6 | 30/09/21 5:49 pm | 01/10/21 9:15 am |
| T2 | Closed forest | 55°38'19"N 11°22'6"E | 1 | 28/09/21 8:01 am | 28/09/21 5:55 pm |
| | | | 2 | 28/09/21 6:07 pm | 29/09/21 7:43 am |
| | | | 3 | 29/09/21 7:55 am | 29/09/21 5:55 pm |
| | | | 4 | 29/09/21 6:05 pm | 30/09/21 7:43 am |
| | | | 5 | 30/09/21 7:54 am | 30/09/21 5:51 pm |
| | | | 6 | 30/09/21 6:05 pm | 01/10/21 9:21 am |
| T1 | Open forest | 55°38'20"N | 1 | 28/09/21 | 28/09/21 |

| | | | | | |
|---|---|---|---|---|---|
| | | 11°22'9"E | | 08:37 am | 6:23 pm |
| | | | 2 | 28/09/21<br>6:36 pm | 29/09/21<br>8:10 am |
| | | | 3 | 29/09/21<br>8:19 am | 29/09/21<br>6:21 pm |
| | | | 4 | 29/09/21<br>6:29 pm | 30/09/21<br>8:09 am |
| | | | 5 | 30/09/21<br>8:18 am | 30/09/21<br>6:19 pm |
| | | | 6 | 30/09/21<br>6:31 pm | 01/10/21<br>9:35 am |
| T2 | Open forest | 55°38'20"N<br>11°22'6"E | 1 | 28/09/21<br>8:20 am | 28/09/21<br>6:08 pm |
| | | | 2 | 28/09/21<br>6:21 pm | 29/09/21<br>7:57 am |
| | | | 3 | 29/09/21<br>8:08 am | 29/09/21<br>6:07 pm |
| | | | 4 | 29/09/21<br>6:17 pm | 30/09/21<br>7:57 am |
| | | | 5 | 30/09/21<br>8:06 am | 30/09/21<br>18:06 pm |
| | | | 6 | 30/09/21<br>6:17 pm | 01/10/21<br>9:26 am |
| T1 | River side | 55°38'22"N<br>11°22'9"E | 1 | 28/09/21<br>8:51 am | 28/09/21<br>6:38 pm |
| | | | 2 | 28/09/21<br>6:51 pm | 29/09/21<br>8:21 am |
| | | | 3 | 29/09/21<br>8:37 am | 29/09/21<br>6:33 |
| | | | 4 | 29/09/21<br>6:49 pm | 30/09/21<br>8:20 am |
| | | | 5 | 30/09/21<br>8:29 am | 30/09/21<br>6:34 pm |
| | | | 6 | 30/09/21<br>6:46 pm | 01/10/21<br>9:35 am |
| T2 | River side | 55°38'22"N<br>11°22'6"E | 1 | 28/09/21<br>9:17 am | 28/09/21<br>6:54 pm |
| | | | 2 | 28/09/21 | 29/09/21 |

| | | | | | |
|---|---|---|---|---|---|
| | | | | 7:08 pm | 8:42 am |
| | | | 3 | 29/09/21<br>8:57 am | 29/09/21<br>6:53 pm |
| | | | 4 | 29/09/21<br>7:04 pm | 30/09/21<br>8:35 am |
| | | | 5 | 30/09/21<br>8:46 am | 30/09/21<br>6:49 pm |
| | | | 6 | 30/09/21<br>7:02 pm | 01/10/21<br>9:55 am |

**Supplementary methods**

*DNA extraction*

For an initial screening, one of the field replicates from six samples were extracted using the DNeasy Blood & Tissue Kit (QIAGEN). However, a qPCR screening using the 12S vertebrate primer showed the presence of inhibitors and therefore, the eluted DNA was cleaned using the OneStep PCR Inhibitor Removal kit (Zymo Research). To compare the results of this method with another extraction kit, the DNA from the second field replicates the same six samples were extracted using the Power Soil Pro kit (QIAGEN), following the manufacturer's instructions. The differences between extraction kits were screened using a second qPCR, which indicated that the DNA extracted using the combination of the Blood and Tissue and the OneStep PCR Inhibitor removal showed the best results. Therefore the rest of the samples were extracted using those kits.
